# Targeted memory reactivation of a serial reaction time task in SWS, but not REM, preferentially benefits the non-dominant hand

**DOI:** 10.1101/2020.11.17.381913

**Authors:** Anne C. M. Koopman, Mahmoud E. A. Abdellahi, Suliman Belal, Martyna Rakowska, Alun Metcalf, Monika Śledziowska, Thomas Hunter, Penny Lewis

## Abstract

Targeted memory reactivation (TMR) is a technique by which sounds paired with learned information can be used to cue neural reactivation of that information during sleep. While TMR in slow-wave sleep (SWS) has been shown to strengthen procedural memories, it is unclear whether TMR in rapid eye movement (REM) sleep, a state strongly associated with motor consolidation, provides equivalent benefit. Furthermore, it is unclear whether this technique influences dominant and non-dominant hands equally. We applied TMR of a two-handed serial reaction time task (SRTT) during either SWS or REM in thirty-two human right handed adults (sixteen female) to examine the impact of stimulation in each sleep stage on right (dominant) and left hands. While TMR in SWS led to strong benefits in reaction times and sequence-specific skill, equivalent cueing in REM led to no benefit at all, suggesting that reactivation in this sleep stage is not important for the SRTT. Event-related potentials elicited by TMR cues for left and right hand movements differed significantly in REM, but not SWS, showing that these cues are at least processed in REM. Interestingly, TMR benefits were apparent only in the non-dominant hand, potentially due to the weaker performance measured in this hand at the outset. Overall, these findings suggest that memory replay in SWS, but not REM, is important for consolidation of the SRTT, and TMR-cued consolidation is stronger in the non-dominant hand.

**Significance statement:** Targeted memory reactivation (TMR) in sleep leads to memory consolidation, but many aspects of this process remain to be understood. We used TMR of a bimanual serial reaction time task to show that behavioural benefit is only observed after stimulation in SWS, even though electrophysiology shows that the TMR cues are processed in REM. Importantly, TMR selectively benefitted the non-dominant hand. These findings suggest that TMR in REM does not benefit this serial reaction time task, and that TMR in SWS preferentially consolidates weaker memory traces relating to the non-dominant hand.

## Introduction

Memories consolidate across sleep (Diekelmann and Born, 2010; Rasch and Born, 2013), and this is facilitated by offline reactivation in which task-related brain activity is reinstated during sleep (Wilson and McNaughton, 1994; Skaggs and McNaughton, 1996). Targeted memory reactivation (TMR) can be used to influence memory consolidation by biasing memory reactivation. This can be achieved by presenting sounds or smells that were previously linked to items learned in wake during subsequent sleep. TMR has been shown to trigger memory reactivation (Belal et al., 2018; Schreiner et al., 2018; Shanahan et al., 2018) and to influence memory performance after sleep, for example by improving episodic (Rasch et al., 2007; Rudoy et al., 2009; Cellini and Cappuzo, 2018) and procedural skill consolidation (Antony et al., 2012; Schönauer et al., 2014).

TMR is often applied during non-REM sleep stages such as slow-wave sleep (SWS) or Stage 2 (e.g. Antony et al., 2018; Cairney et al., 2014; Fuentemilla et al., 2013; Hauner et al., 2013; Rasch et al., 2007; Rudoy et al., 2009). Nevertheless, early studies show that tasks with a procedural memory component benefit from rapid eye movement (REM) sleep (Smith, 1993, 1995, 2001; Karni et al., 1994; Smith and Smith, 2003) and brain regions involved in the serial reaction time task (SRTT) are reactivated during REM (Maquet et al., 2000; Peigneux et al., 2003). Furthermore, rodent studies have clearly identified replay in REM (Poe et al., 2000; Louie and Wilson, 2001; Booth and Poe, 2006; Howe et al., 2019), while human work has suggested that REM replay may be stronger than replay in other sleep stages (Schönauer et al., 2017). The SRTT is a reaction time task which has already been shown to be sensitive to TMR in SWS (Cousins et al., 2014, 2016). In a magnetic resonance imaging study, we showed that widespread plasticity in the motor system in response to TMR of this task in SWS is mediated by time spent in REM. Nevertheless, the influence of TMR in REM on this task has never been examined. Given the literature, we set out to compare the impact of TMR in REM and SWS on the SRTT.

Studies of sleep-dependent consolidation in motor skills tend to focus on the non-dominant hand (Walker et al., 2002, 2003a, 2003b, 2005; Korman et al., 2003, 2007; Spencer et al., 2006). This is also true for TMR studies of procedural memory (Antony et al., 2012; Cousins et al., 2014, 2016; Schönauer et al., 2014). The non-dominant hand is typically chosen to reduce the influence of pre-existing motor skills (Maquet et al., 2003) and because greater performance gains are possible in this hand (Ridding and Flavel, 2006; Spencer et al., 2006). However, we are unaware of any study comparing the impact of sleep dependent memory consolidation on dominant and non-dominant hands. We therefore set out to examine this within our SRTT. Our right-handed participants performed a two-handed variant of this task which allowed us to examine the impact of TMR on task behaviour in both dominant and non-dominant hands. TMR cueing was applied to separate groups of participants in REM and SWS to allow examination of the importance of sleep stage.

## Materials and Methods

### Participants

Thirty-five healthy right-handed, non-smoking participants were recruited for this study and randomly assigned to either the SWS- or REM-group, with the constraint of gender balance. Three participants were excluded, due to spending <30 minutes in SWS during the experimental night (*n* = 1), experimenter error resulting in TMR during wake (*n* = 1), and no evidence of learning during the pre-sleep training (*n* = 1). Sixteen participants remained in the REM Cued group (8 female, mean age 23.6 years) and sixteen in the SWS Cued group (8 female, mean age 23.1 years).

All participants had normal or corrected-to-normal vision, normal hearing, and no history of physical, psychological, neurological, or sleep disorders. Responses in a pre-screening questionnaire reported no stressful life events, a generally regular sleep-wake rhythm in the month before the study, and no regular night work or cross-continental travel in the two months before the study. Participants were not taking any psychologically active medication or substances and agreed to abstain from alcohol and caffeine in the 24 and 12 hours prior to the start of the study, respectively. Subjects also agreed not to nap or participate in extreme physical exercise during the experiment, starting 12 hours before their first arrival in the lab. This study was approved by the School of Psychology, Cardiff University Research Ethics Committee, and all participants gave written informed consent.

### Experimental tasks and design

#### Design

The study consisted of an adaptation night, which allowed participants to get used to sleeping in the lab with electrodes, and an experimental night during which they performed the behavioural tasks. The study design is outlined in Figure 1. There was at least one but no more than three nights between the adaptation and experimental nights.

**Figure 1.**
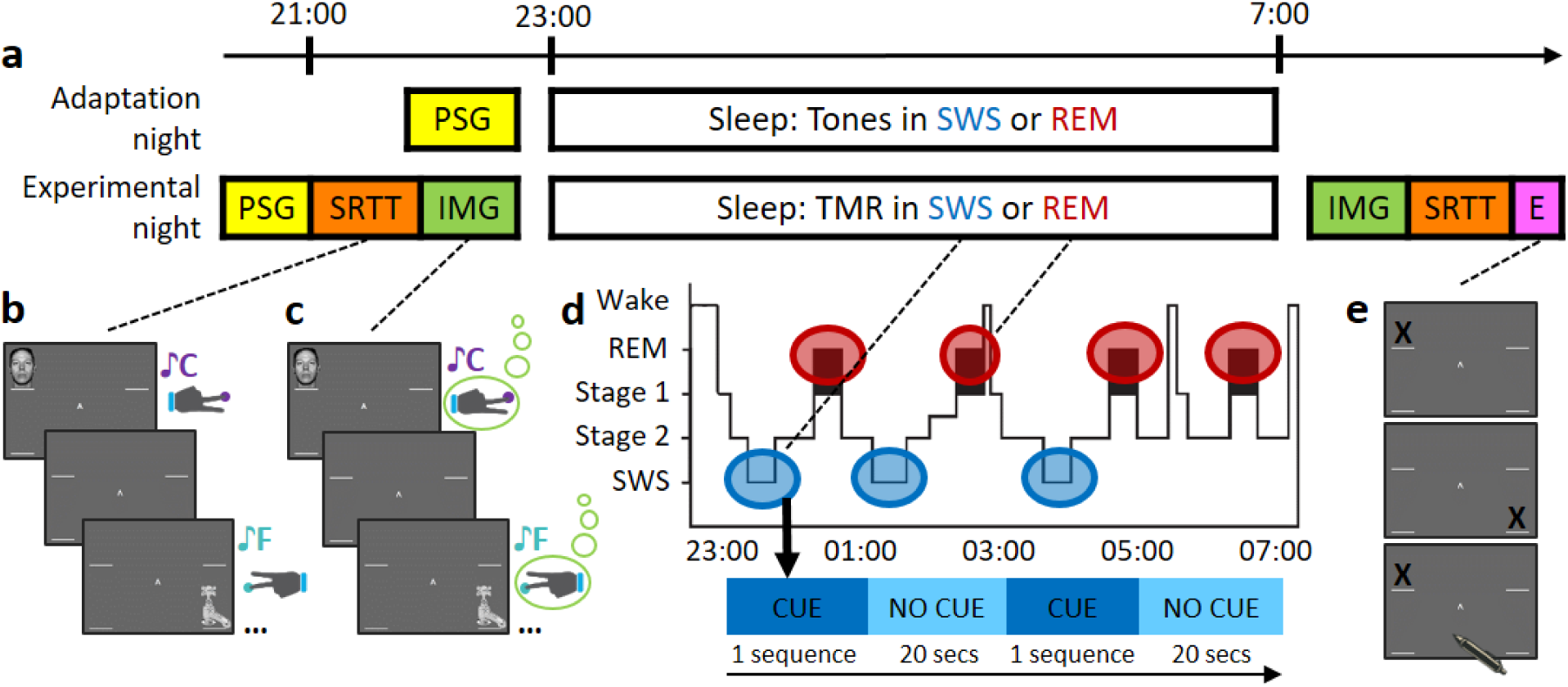
Experimental procedures. **a)** The experiment consisted of an adaptation and an experimental night. During the adaptation night, participants were wired-up for EEG and, while they slept, tones were played as outlined in d). During the experimental night, participants were wired-up, after which they completed the SRTT and IMG tasks as outlined in b) and c), respectively. Then, participants went to sleep and TMR was carried out as described in d). After waking up, participants completed the IMG and SRTT again, and finally the explicit recall task (E) which is described in e). **b)** Serial Reaction Time Task (SRTT). Images were presented in two different sequences. Each image was accompanied by a specific pure tone (different for each sequence) and required a specific button press. **c)** Motor imagery task (IMG). Participants viewed and heard the same sequences again, but this time were instructed to only imagine pressing the buttons. **d)** Schematic representation of the TMR protocol. Reactivation took place in either SWS (blue bubbles) or REM sleep (red bubbles). One sequence was played as long as participants were in the relevant sleep stage, with a 20 second pause between repetitions. **e)** Explicit recall task. Participants marked the order of each sequence on paper.

Upon arrival for the adaptation night, around 9:30pm, participants changed into their sleepwear. At this time, their ability to use internal visual and kinaesthetic imagery was measured with a shortened version of the Movement Imagery Questionnaire-3 (MIQ-3; Williams et al., 2012) and handedness was assessed with a short version of the Edinburgh Handedness Inventory (EHI; Veale, 2014). They were then fitted for polysomnography (PSG) recording. Subjects’ alertness was assessed by the Karolinska Sleepiness Scale (KSS; Åkerstedt & Gillberg, 1990) and the Stanford Sleepiness Scale (SSS; Hoddes, Zarcone, Smythe, Phillips, & Dement, 1973) before going to bed around 11-11:30pm. During this night, we played the tones that the participants would later (during the experimental training) learn to associate with one of the learned sequences. As these tones had not yet been associated with a memory, playing them during the adaptation night meant that they could be used as a control for analyses looking at the neural signature of memory reactivation. After 7-8 hours of sleep, participants were awakened. They then rated the sleep quality of the night with an adapted and translated version of a German sleep quality questionnaire (SQQ; Görtelmeyer, 1985), and completed the KSS and SSS again. After removing the electrodes, participants were offered the opportunity to shower before leaving the lab.

On the experimental night, participants arrived around 7:30pm and then changed into their sleepwear. They completed the Pittsburgh Sleep Quality Index (PSQI; Buysse, Reynolds, Monk, Berman, & Kupfer, 1989) to report their sleep quality over the past month and were then fitted for PSG recording. Participants performed the serial reaction time task (SRTT; 50-60 minutes) and the motor imagery task (IMG; 30 minutes). Before each task, the KSS and SSS were completed to measure alertness. Participants were ready for bed around 11pm. During the night, the tones that had been played during the adaptation night, and again during one of the learned sequences, were replayed. The sleep stage of cueing was the same in experimental and adaptation nights (SWS or REM, depending on the group the participant was in). As these tones were now associated with the SRTT, we expected that cueing them would trigger reactivation of this association (a memory) (Cousins et al., 2014, 2016; Belal et al., 2018). After 7-8 hours of sleep, participants were awakened and allowed at least 20 minutes to overcome sleep inertia. During this time, participants were given the opportunity to eat and drink something before completing the sleep quality questionnaire. Participants then completed the same tasks again in reverse order (IMG first, SRTT second), each preceded by the KSS and SSS. Finally, participants completed an explicit sequence memory test, by marking the sequence order on a printout containing pictures of the (empty) screen arranged vertically (Figure 1e). This was done for both sequences; the order was counterbalanced across participants. Tasks were presented on a computer screen with resolution 1024 x 768 pixels and using Matlab 6.5 (The MathWorks Inc., Natick, MA, 2000) and Cogent 2000 (Functional Imaging Laboratory, Institute for Cognitive Neuroscience, University College London). The tones were played through noise-cancelling headphones (Sony MDR-ZX110NA) during the tasks and through speakers (Dell A225) during sleep.

#### Serial Reaction Time Task (SRTT)

The main task was a serial reaction time task (SRTT; adapted from Cousins et al., 2014; see Figure 1b) which contained two sequences of twelve items that were learned in interleaved blocks. The sequences – A: 1 2 1 4 2 3 4 1 3 2 4 3 and B: 2 4 3 2 3 1 4 2 3 1 4 1 – had been matched for learning difficulty, did not share strings of more than four items, and both contained each item three times. The blocks were interleaved so that a block of the same sequence was presented no more than twice in a row, and each block contained three repetitions of a sequence. There were 24 blocks of each sequence (48 blocks in total), and each block was followed by a pause of 15 seconds wherein feedback on reaction time (RT) and error-rate was presented. The pause could be extended by the participants if they wanted. Participants were aware that there were two twelve-item sequences and each sequence was indicated with ‘A’ or ‘B’ appearing centrally on the screen, but participants were not asked to learn the sequences explicitly. Counterbalancing across participants determined whether sequence A or B was the first block, and which of the sequences was cued during sleep.

Each sequence was paired with a group of pure musical tones, either low tones within the 4^th^ octave (C/D/E/F) or high tones within the 5^th^ octave (A/B/C#/D). These tone groups were counterbalanced across sequences. For each trial, a 200 ms tone was played, and at the same time a visual cue appeared in one of the corners of the screen. The location indicated which key on the keyboard needed to be pressed as quickly and accurately as possible: 1 – top left corner = left Shift; 2 – bottom left corner = left Ctrl; 3 – top right corner = up arrow; 4 – bottom right corner = down arrow. Participants were instructed to keep individual fingers of their left and right hand on the left and right response keys, respectively. Visual cues were neutral objects or faces, used in previous studies (Cousins et al., 2014, 2016). Stimuli appeared in the same position for each sequence (1 = male face, 2 = lamp, 3 = female face, 4 = water tap) and participants were instructed that the nature of the cues (objects/faces) was irrelevant. Visual cues stayed on the screen until the correct key was pressed, after which an 880 ms inter-trial interval followed.

After the 48 blocks of sequences A and B, participants performed four more blocks that contained semi-random sequences which followed only the rule that no item was presented twice in a row. They contained the same visual stimuli and an ‘R’ displayed centrally on the screen. Two of these blocks were paired with the tone group of one sequence (cued in sleep), and the other two were paired with the tone group of the other sequence (not cued).

#### Motor Imagery Task (IMG)

After completion of the SRTT, participants were asked to do the same task again, but were instructed to only imagine pressing the buttons (Figure 1c). This task consisted of 30 interleaved blocks (15 of each sequence), presented in the same order as during the SRTT. Again, each trial consisted of a 200 ms tone and a visual stimulus, the latter being shown for 880 ms and followed by a 270 ms inter-trial interval. There were no random blocks during this motor imagery task (IMG) and no performance feedback was presented during the pause between blocks.

#### TMR during REM and SWS

Cueing was started when participants – depending on their assigned group – were in stable REM sleep or SWS (fitting standard AASM criteria for Stage R or N3). Tones were presented as often as possible, with a pause of 1500 ms between tones. One repetition of the sequence was presented, alternated by a 20 second break. Figure 1d shows a schematic representation of the TMR protocol. Cueing was paused immediately when participants showed any sign of arousal or when they left the relevant sleep stage. When a return to stable SWS/REM sleep was apparent, cueing was continued.

In the REM group, on average 984 sounds were played in the adaptation night (range 540-1404) and 1177 sounds in the experimental night (range 768-1718). In the SWS group, an average of 1159 sounds were played in the adaptation night (range 338-1836) and 1272 sounds in the experimental night (range 978-1908). There was no significant difference between the groups during the adaptation night (t(30) = −1.37; p = 0.181) or the experimental night (t(30) = −0.97; p = 0.342), as tested with two independent samples t-tests.

### PSG data acquisition and analysis

Twenty-one electrodes were placed on the scalp and face of the participants following the 10-20 system. On the scalp, these were at 13 standard locations: Fz, Cz, Pz, F3, F4, C5, CP3, C6, CP4, P7, P8, O1, and O2, and they were referenced to the mean of the left and right mastoid electrodes. Further electrodes used were the left and right EOG, three EMG electrodes on the chin, and the ground electrode on the forehead. The impedance was <5 kΩ for each scalp electrode, and <10 k*Ω* for each face electrode. Recordings were made with an Embla N7000 amplifier and RemLogic 1.1 PSG Software (Natus Medical Incorporated). PSG recordings were manually scored by two trained and independent sleep scorers according to the standard AASM criteria (Berry et al., 2015). Both scorers were blind to the periods cueing occurred.

### Electrophysiological analysis

The EEG data that were collected using the methods described above were further analysed with MATLAB (version R2016b) and the FieldTrip Toolbox (version 20/08/2019, Oostenveld, Fries, Maris, & Schoffelen, 2011). First, the raw data was coupled to the identity of the sounds played during sleep. Then, these marked continuous data were filtered between 0.1 and 30 Hz and coupled to the sleep scoring. This allowed for the removal of any trials where the tones were played during the wrong sleep stage or during an arousal. Subsequently, the continuous data were segmented into trials starting 1 second before sound onset and ending 3 seconds after sound onset. Trials that took place during the wrong stage or during an arousal were discarded. In the adaptation night, this resulted in the removal of on average 5.8% and 6.4% of data in the REM and SWS groups, respectively. In the experimental night, this resulted in the removal of on average 4% of data in the REM and 10.7% of data in the SWS group.

Further artifacts were removed in a multi-step procedure. Trials were first re-segmented into smaller trials of −0.5 and +3 seconds around the onset of a sound. Each EEG channel was then looked at separately. A trial was considered an outlier for a given channel if it was more than two standard deviations from the mean on amplitude or variance. If a trial was considered an outlier in more than 25% of channels (i.e. 3 channels), then this trial was rejected. During this procedure, on average 10.5% of trials were removed from the adaptation night data in the REM group, and 11.9% of trials in the SWS group. In the experimental night, this led to the removal of 9.8% of trials in the REM, and 11.7% of trials in the SWS group. Those trials which were considered an outlier in <25% of channels were interpolated based on triangulation of neighbouring channels. Data in the REM group was subsequently analysed with independent component analysis (ICA), to remove eye movement artifacts which can occur during REM. Components identified by the ICA were correlated with the signal from the eye electrodes, and components that were significantly correlated (corrected for multiple comparisons) were removed. In the final artifact rejection step, all channels for each participant were manually inspected. Any channels that showed overall noise were interpolated based on their neighbours, and overly noisy trials still detected during this step were removed.

Event-related potentials (ERPs) were analysed time-locked to TMR cue start. To reduce influence of outliers, we used the median to calculate the ERP of the segments. Grand-averages were baseline-corrected to a baseline window of −1 second until cue onset. This long baseline window was chosen because of the low-frequency nature of SWS. ERPs were expected to occur shortly after cue onset, and therefore statistical analyses focused on the time period between cue onset and 500 ms thereafter. Due to the pronounced difference in electrophysiology between SWS and REM, comparisons between cues (left-versus right-handed cues, and cues during the adaptation versus experimental night) were done separately for the SWS and REM groups. They were performed as paired-samples t-tests and corrected for multiple comparisons using FieldTrip’s nonparametric cluster-based permutation method, using 1000 permutations. Results were considered significant at p <0.05.

We tested for correlations between the ERP difference between the adaptation and experimental night with behavioural improvement in the experimental night. To this end, we looked at the 60 ms surrounding the early evoked peak in each group, averaged these intervals across each night, and computed the difference by subtracting the adaptation from the experimental night. This difference value was then correlated with overnight behavioural improvement.

### Spindle analysis

Previous studies using similar tasks have indicated a relationship between sleep spindles (short bursts of activity in the 11-16 Hz range) over central or frontal electrodes and behavioural outcomes (Nishida and Walker, 2007; Antony et al., 2012; Cousins et al., 2014). Therefore, we examined spindles in the SWS group, using a counting algorithm based on one used by Antony and colleagues (2018). In short, the raw EEG was filtered in sigma band as identified in the paper by Antony et al. (11-16 Hz), and root-mean-square (RMS) values were calculated using a sliding 200 ms window. Segments that fit threshold criteria were selected. To fit threshold, a spindle had to be during stage N3 (SWS), be between 0.3 and 2.5 seconds long, and have at least 5 oscillations in that period. Spindle identity was then further confirmed by using time-frequency information to detect whether increased spindle-band power indeed took place during the selected segments (Purcell et al., 2017; Navarrete et al., 2020).

After detection, any spindles that fell partly or wholly during a previously visually marked arousal were removed. The remaining spindles were divided into those taking place during TMR, and those taking place during baseline SWS. If a spindle started up to 1.65 seconds after tone onset, it was considered to occur during TMR. This interval was chosen because subsequent tones in the sequence, when a sequence was played without pauses, would be a maximum of 1.65 seconds apart. If the start of a spindle fell outside of this interval, it was considered to occur during baseline. Spindle density was calculated as the amount of ‘within TMR’ spindles divided by the amount of cues played during the night. We then computed spindle laterality in the left hemisphere by subtracting spindle density in the right from the left hemisphere. To obtain spindle laterality in the right hemisphere, finally, we subtracted spindle density in the left from that in the right hemisphere. Because Cousins and colleagues (2014) found an effect of spindle laterality in central electrodes on procedural skill improvement in the SRTT, our analyses focused on electrodes C5 & CP3 (left hemisphere) and C6 & CP4 (right hemisphere). Notably, because our participants used both hands, while the participants in prior studies showing a relationship between spindle laterality and finger tapping improvement across sleep have used just one hand (Walker et al., 2002; Antony et al., 2012; Cousins et al., 2014, 2016), we did not necessarily expect to find a relationship.

### Behavioural analysis

Performance on the SRTT was measured by the reaction time (RT) per block. Following the method used by Cousins et al. (Cousins et al., 2014, 2016), any trials with an RT of more than 1000 ms were excluded from analyses, while trials with incorrect button presses prior to the correct ones were not excluded. Because we were interested in differences between the dominant and non-dominant hands, as well as the overall performance, the data were analysed in three different ways: Both Hands (BH), Left Hand (LH), and Right Hand (RH).

Subjects were excluded if (1) their RT performance before sleep was >2 SDs from the group mean (*n* = 1 in the LH dataset), (2) there was a >2 SD disparity between the group mean RT for the two sequences before sleep (*n* = 1 in the BH dataset, *n* = 2 in the LH dataset, and *n* = 2 in the RH dataset), or (3) they exhibited a positive slope of the learning curve before sleep, i.e. they did not show any learning during training (*n* = 1 for all datasets). Thus, thirty participants remained in the BH dataset (*n* = 15 for the REM group, and *n* = 15 for the SWS group), twenty-eight in the LH dataset (*n* = 15 for the REM group, and *n* = 13 for the SWS group), and twenty-nine in the RH dataset (*n* = 15 for the REM group, and *n* = 14 for the SWS group).

RTs were divided into those for the cued and those for the uncued sequence. Performance on the last four blocks before sleep was considered to represent pre-sleep ability, and this was subtracted from the random blocks to remove the effects of increased sensorimotor mapping ability. The resulting variable can thus be called sequence-specific skill. Sequence-specific improvement was then calculated for each sequence by subtracting the pre-sleep sequence-specific skill from the post-sleep sequence-specific skill (i.e. the random blocks after sleep minus the first 4 blocks after sleep). Higher values on this value thus indicate more improvement.

We used analyses of variance (ANOVAs) to determine the effects of time (before/after sleep) and sequence (cued/uncued) within each group (REM/SWS) separately. As follow-up, we used t-tests or Wilcoxon singed-rank tests (whenever the Shapiro-Wilk test indicated non-normal distribution). Relationships between behavioural measures and features of the EEG and sleep were assessed with Pearson’s correlations, or Spearman’s Rho in the case of non-normal distributions. All statistical tests were 2-tailed and considered significant for p < 0.05. Analyses were conducted in R (version 3.6.3, R Core Team, 2020). We included measures of effect size: generalised eta squared η^2^_*G*_ for ANOVA as calculated with the “afex” R package (Olejnik and Algina, 2003; Bakeman, 2005; Lakens, 2013; Singmann et al., 2020), and *r* for Wilcoxon tests as calculated with the “rcompanion” R package (Fritz et al., 2012; Mangiafico, 2020).

## Results

### Sleep parameters

Sleep scoring confirmed that the vast majority of all TMR sounds were played during the correct sleep stage. No sounds were played in the opposite sleep stage (i.e. during SWS for the REM group, and vice versa). Sleep parameters did not differ between groups, with the notable exception of Stage 2, which was longer in the SWS group during both the adaptation night (F(1,30) = 6.2; p = 0.018) and the experimental night (F(1,30) = 6.1; p = 0.020). A summary of the time spent in sleep stages can be found in Table 1.

**Table 1.**
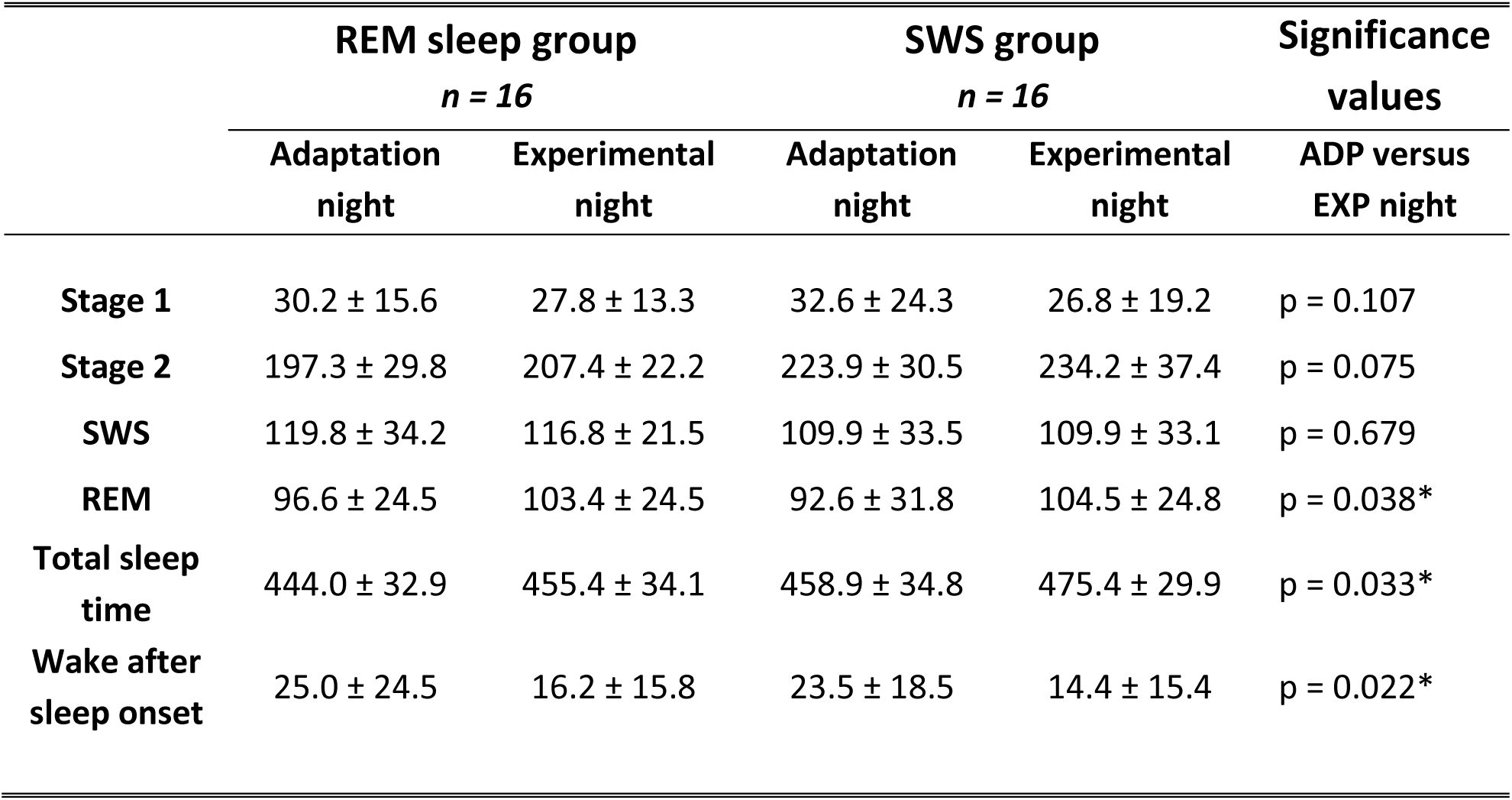
Average minutes spent in sleep stages (± standard deviation). * = significant at the α = 0.05 level.

We combined the groups and conducted paired t-tests to see whether sleep was better on the second night spent in the lab. Participants slept significantly longer in the experimental compared to the adaptation night (t(31) = −2.23; p = 0.033, see Table 1). Participants also spent more time in REM in the experimental night than the adaptation night (t(31) = −2.16; p = 0.038), but the difference in Stage 2 sleep did not reach significance (t(31) = −1.84; p = 0.075). There was also no difference in SWS (t(31) = 0.41; p = 0.679) or Stage 1 sleep (t(31) = 1.66; p = 0.107). On the other hand, participants did wake up more during the adaptation night compared to the experimental night (t(31) = 2.42; p = 0.022), which may contribute to the difference in total sleep time across the nights.

We also evaluated self-rated quality of sleep across nights, and self-rated positive feeling on the mornings after waking up in the lab. In terms of sleep quality, on average this was rated higher after the experimental compared to the adaptation night (t(31) = −2.32; p = 0.027). However, when evaluating the ratings of how participants felt in the morning, we found no difference between the nights (t(31) = 0.25; p = 0.808).

### Behaviour: Both Hands

We first evaluated behavioural results for both hands combined. At the end of the pre-sleep training, RTs were significantly faster for trials within a sequence compared to trials within the random blocks, confirming learning of both sequences in both the REM and the SWS groups (all p < 0.001). Importantly, during pre-sleep baseline, RTs did not differ between the cued and uncued sequences for either sequence or for the random blocks (lowest p: F(1,28) = 1.22; p = 0.280).

An ANOVA with factors time (before and after sleep) and TMR (cued or uncued sequence) examined sequence-specific skill change in SWS and REM groups. In SWS, as expected, there was a main effect of time (F(1,14) = 11.35; p = 0.005; η_G_^2^ = 0.108), with faster performance after sleep. There was also a time x TMR interaction (F(1,14) = 5.01; p = 0.042; η_G_^2^ = 0.015), with more improvement shown on the cued versus the uncued sequence, as determined with a Wilcoxon signed-rank test (V = 84; p = 0.049; r = 0.529; pictured in Figure 3e). The main effect of TMR across pre and post-sleep sessions was not significant (F(1,14) = 0.33; p = 0.574). In the REM group, our ANOVA showed a main effect of time (F(1,14) = 44.77; p < 0.001; η_G_^2^ = 0.137), but no other main effects or interaction (lowest p: F(1,14) = 0.37; p = 0.553), Figure 2e.

**Figure 2.**
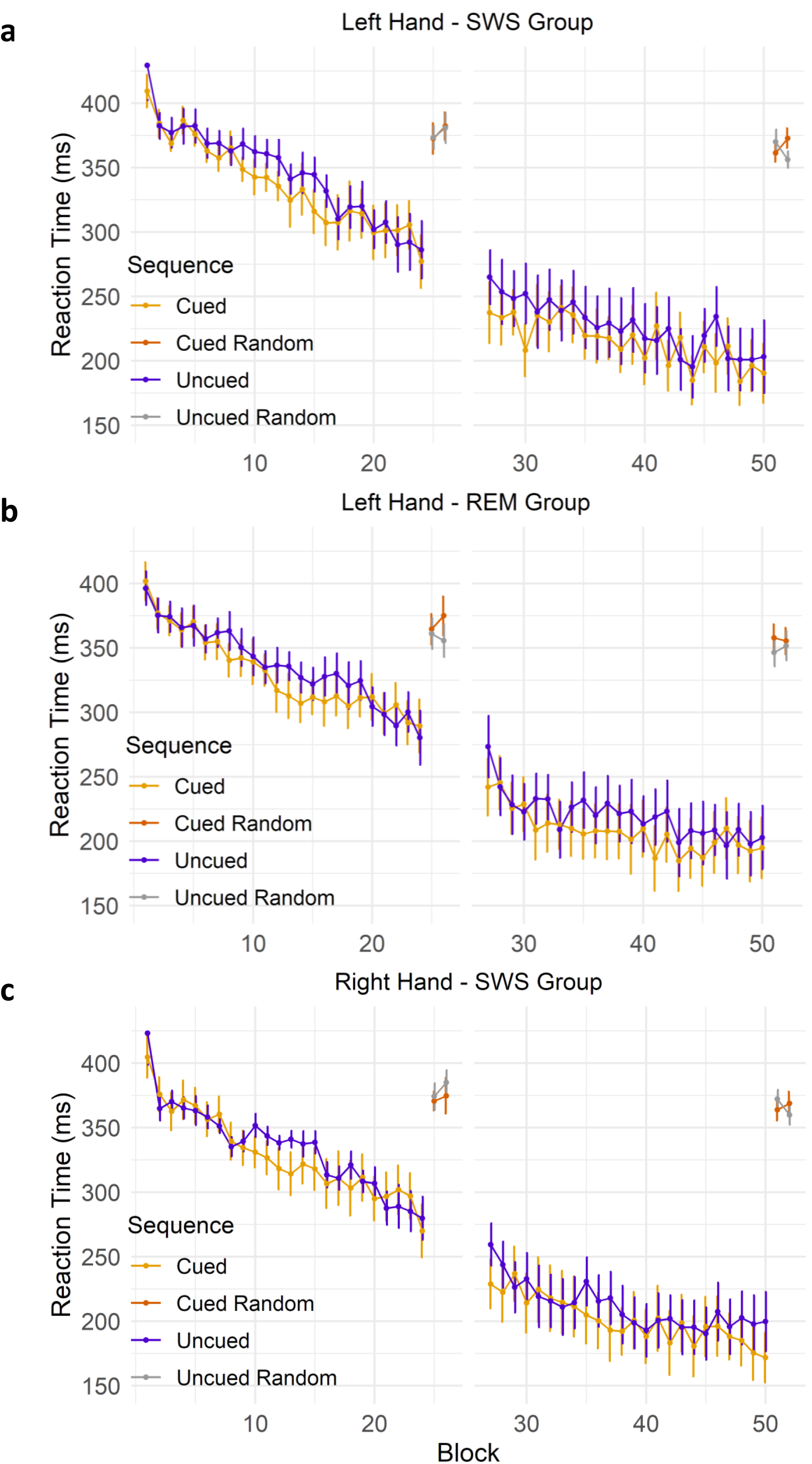
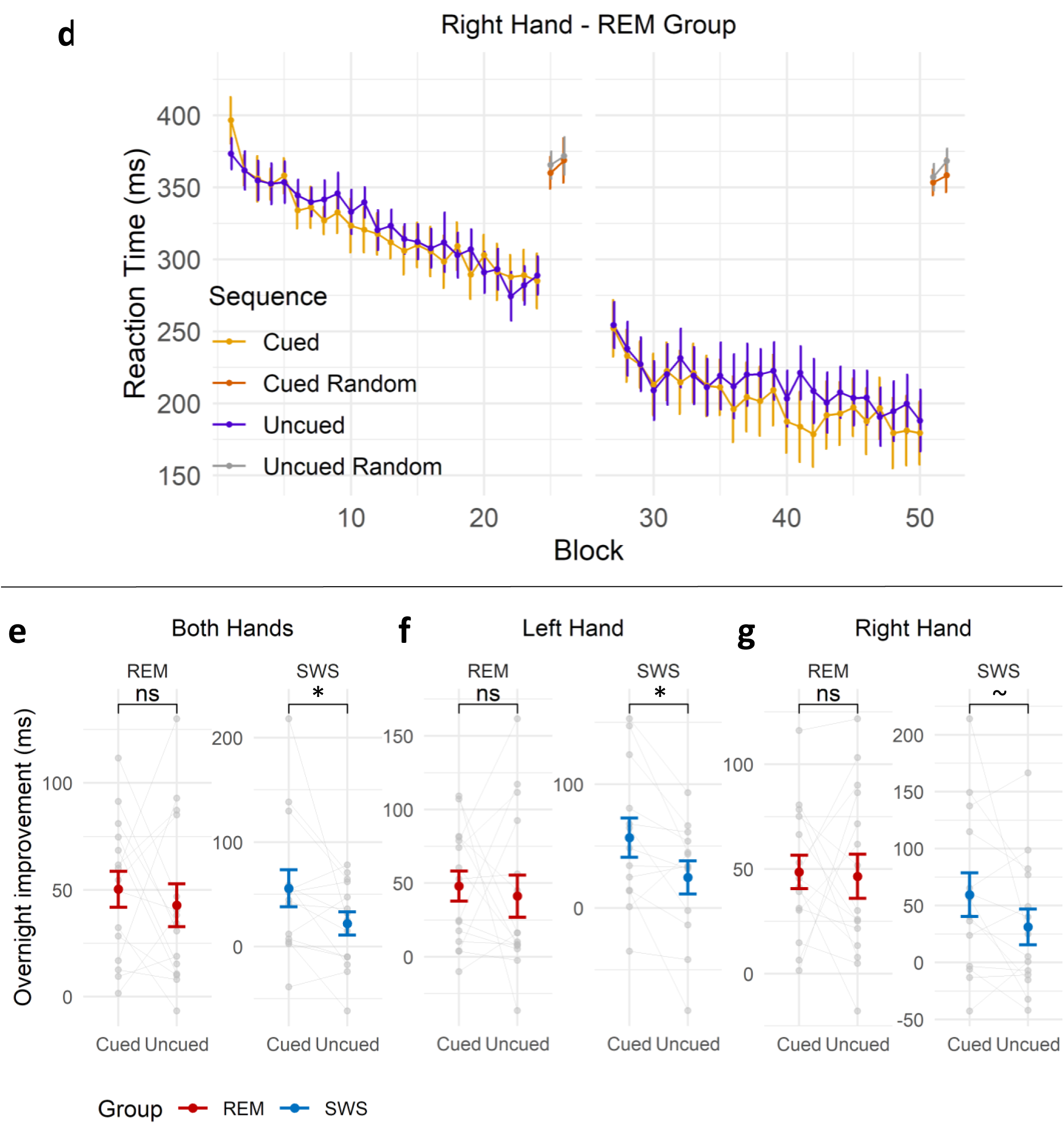
Behavioural results. **a)** SRTT performance across all blocks of learning, for the SWS group in the left hand. **b)** SRTT performance across all blocks of learning, for the REM group in the left hand. **c)** SRTT performance across all blocks of learning, for the SWS group in the right hand. **d)** SRTT performance across all blocks of learning, for the REM group in the right hand. **e)** SRTT overnight sequence-specific improvement in both hands was significantly better for the cued than uncued sequence in the SWS group only (V = 84; p = 0.049). **f)** SRTT overnight sequence-specific improvement in the left hand was greater for the cued than uncued sequence in the SWS group only (V = 79; p = 0.017). **g)** SRTT overnight sequence-specific improvement in the right hand did not differ between cued and uncued sequence in either group (Difference in the SWS group: t(13) = 1.73; p = 0.108). Data are presented as mean ± SEM.

**Figure 3.**
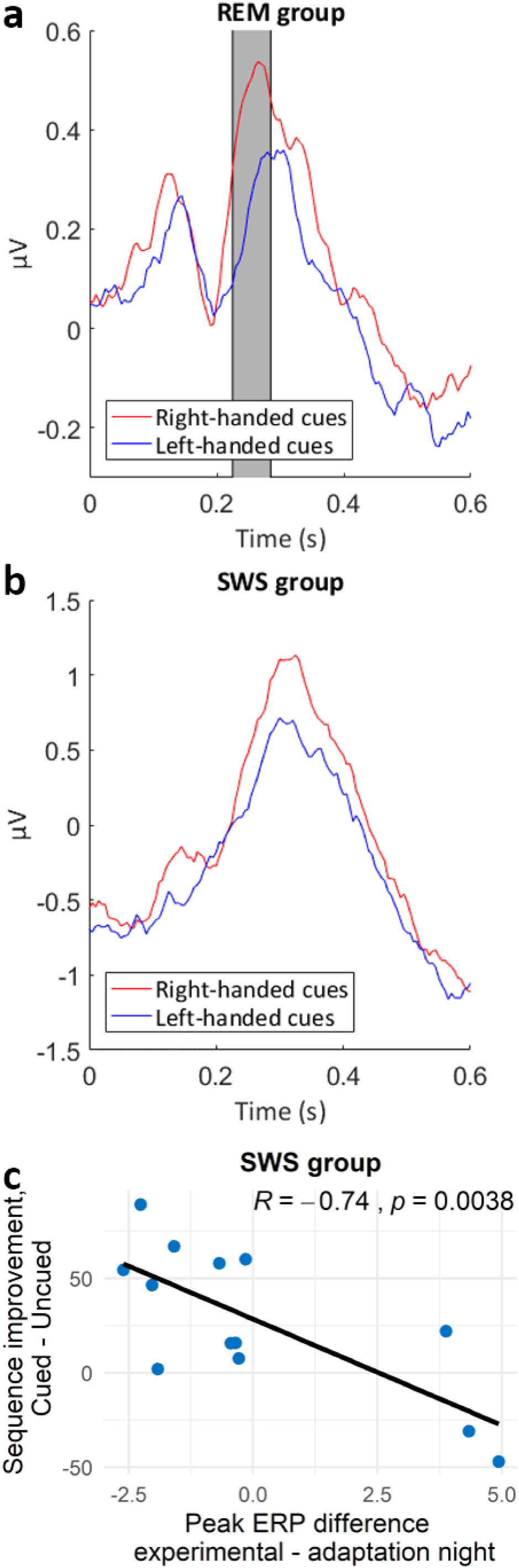
ERPs to left- and right-handed cues over all EEG channels during the experimental night **a)** In the REM group. Grey rectangle indicates when the difference between these cues is significant (p < 0.05; corrected for multiple comparisons). **b)** In the SWS group. **c)** Significant correlation between the overnight sequence improvement cue benefit in the left hand and the ERP difference of the experimental versus the adaptation night at the first peak after the tone (p = 0.015 after correction for multiple comparisons).

### Behaviour: Left and Right Hand

To investigate whether TMR influenced each hand similarly, we separated trials into those where responses were made with the left (non-dominant) or right (dominant) hand. At the end of the pre-sleep training, performance of both cued and uncued sequences was significantly faster than performance on the random blocks (all p < 0.001) in both right hand (RH) and left hand (LH). This confirms learning in both the LH and RH before sleep. Before sleep RTs in either hand did not differ between the cued and uncued sequences, nor between random blocks before sleep (lowest p: F(1,27) = 1.93; p = 0.176), see Figure 2a-d.

We analysed sequence-specific skill change in the REM and SWS groups separately for RH and LH using an ANOVA with within-participant factors time (before and after sleep) and TMR (cued or uncued sequence). Starting with the LH, in the SWS group, this showed a main effect of time (F(1,12) = 9.09; p = 0.011; η_G_^2^ = 0.092) and a time x TMR interaction (F(1,12) = 7.31; p = 0.019; η_G_^2^ = 0.015), showing that TMR had an impact on post-sleep performance, but not on pre-sleep performance. In keeping with this, a Wilcoxon signed-rank test (V = 79; p = 0.017; r = 0.649) showed greater overnight improvement on the cued than the uncued sequence, Figure 2f. As expected, the main effect of TMR (across pre- and post-sleep sessions) was not significant (F1,12) = 1.29; p = 0.279; η_G_^2^ = 0.011). In the REM group, LH showed a main effect of time (F(1,14) = 25.36; p < 0.001; η_G_^2^ = 0.110), but no other effects (lowest p: F(1,14) = 0.80; p = 0.387).

Turning to the RH, this showed no cueing-related improvement in neither the SWS nor REM group. In the SWS group the main effect of time was significant (F(1,13) = 8.63; p = 0.012; η_G_^2^ = 0.113), but there were no other main effects or interaction (lowest p: F(1,13) = 2.98, p = 0.108; η_G_^2^ = 0.012; for the time x TMR interaction). In REM, there was a main effect of time (F(1,14) = 44.47; p < 0.001; η_G_^2^ = 0.144), but no other main effects or interaction (lowest p: F(1,14) = 1.77, p = 0.205). Figure 2g shows the sequence-specific improvement for the cued and uncued sequences per group in the RH.

Given the differences in TMR efficacy in right and left hand, we were interested to know whether either hand showed weaker performance from the outset. We conducted a mixed ANOVA with within-participant factor hand (left or right) and between-participant factor group (REM or SWS) on sequence-specific skill in the last four blocks before sleep. There was a trend towards an effect of hand (F(1,24) = 3.88, p = 0.060; η_G_^2^ = 0.011), with the left hand showing weaker sequence-specific skill than the right hand. There were no other main effects or interactions (lowest p: F(1,24) = 0.35, p = 0.558).

### Behaviour: Reaction Time

Because effects related to handedness might be clearer in reaction times than in sequence-specific skill, we also analysed RT. We first looked at RT in both hands combined. An ANOVA with factors Time (before and after sleep) and TMR (cued or uncued sequence) examined RT change in SWS and REM groups. The time x TMR interaction, our effect of interest, was not significant in neither the SWS (F(1,14) = 3.02; p = 0.104; η_G_^2^ = 0.007) nor the REM group (F(1,14) = 0.68; p = 0.424; η_G_^2^ = 0.002).

In contrast, when looking at the left hand only, there was a significant interaction between time and TMR in the SWS group (F(,12) = 6.41; p = 0.026; η ^2^ = 0.010), with the cued sequence showing more overnight reaction time improvement than the uncued sequence. This interaction was not significant in the REM group (F(1,14) = 0.49; p = 0.494; η_G_^2^ = 0.002). In the right hand, there was no significant time x TMR interaction in the SWS group (F(1,13) = 3.06; p = 0.104; η_G_^2^ = 0.007) nor the REM group (F(1,14) = 0.16; p = 0.694; η_G_^2^ < 0.001). In all cases, the main effect of time was significant (highest p = 0.001) and the main effect of TMR was not significant (lowest p = 0.361).

The RTs show a similar pattern to the sequence-specific skill, where the left hand significantly responded to overnight TMR but the right hand did not. Therefore, we were again interested to know whether either hand showed weaker performance from the outset. Thus, we conducted a mixed ANOVA with within-participant factor hand (left or right), between-participant factor group (REM or SWS), and dependent variable raw RT in the last four blocks before sleep. This showed a main effect of hand (F(1,24) = 15.59, p < 0.001; η_G_^2^ = 0.024), where RT in the left hand was slower than in the right hand. There were no other main effects or interactions (lowest p: F(1,24) = 0.63, p = 0.437). Overall, the left hand showed lower sequence-specific skill and slower raw RT than the right hand before sleep.

### Electrophysiology: Spindle analysis

To determine whether cued sequence advantage related to spindles, we calculated a ‘procedural cueing benefit’ by subtracting cued from uncued sequence RT in the first four blocks after sleep (Cousins et al., 2014). However, there were no significant correlations between spindle density nor laterality and cueing benefit.

### Electrophysiology: Event-related potentials

We examined event-related potentials (ERPs) in response to left- and right-handed TMR cues in sleep for the SWS and REM groups. Interestingly, TMR in REM elicited a stronger response to right-hand compared to left-hand cues when all EEG channels were combined in the experimental night. A cluster-based permutation test on the combined EEG channels and the latency range of 0 to 500 ms post-stimulus showed that this difference was significant between 0.225 and 0.285 seconds after cue onset (p = 0.044; see Figure 3a). Though this difference in right and left hand cues was visible throughout the brain, it was descriptively most pronounced in the left hemisphere (C5 and CP3). In the adaptation night there was no difference between cues associated with the left versus the right hand (lowest p = 0.238). Although TMR in SWS revealed a numerical trend in keeping with the REM results, there were no significant differences in responses to right and left hands in neither experimental (lowest p = 0.447) nor adaptation (lowest p = 0.311) nights. Descriptively, it is interesting to note that TMR cues in this sleep stage showed the slow-oscillatory pattern that is characteristic of slow-wave sleep (see Figure 3b). We further examined ERPs in each hemisphere separately, but this did not reveal any additional information.

Comparison of ERPs in the experimental night and adaptation night, using data from both hands, revealed no significant differences (see supplement). Interestingly however, the extent to which the SWS TMR-related ERP was larger during the experimental than the control night was negatively associated with the overnight LH cue benefit (R = −0.74, p = 0.015; FDR corrected for multiple comparisons; see Figure 3c). To rephrase this, the higher the experimental night peak was, as compared to the adaptation night peak, the less the left hand improved on the cued sequence across the experimental night as compared to the uncued sequence.

## Discussion

In this study, we demonstrate that auditory TMR in SWS, but not REM, was associated with overnight improvement. Furthermore, TMR only benefits behavioural performance in the non-dominant hand, suggesting that the benefit is more marked for memory traces that are weaker at the outset. Despite the literature suggesting that REM sleep is important for motor consolidation, these results support the idea that it is memory reactivation in SWS, not REM, which facilitates overnight improvements in motor sequence performance. Nevertheless, we also found that TMR of right and left handed responses lead to distinct ERPs in REM, but not SWS, showing that REM does process these stimuli at some level.

### TMR in SWS but not REM benefits SRTT consolidation

Early studies of memory consolidation in sleep suggested that REM is critical for motor skills (Smith, 1993, 1995, 2001; Karni et al., 1994; Smith and Smith, 2003). Furthermore, two influential studies showed reactivation of motor networks during REM after a finger tapping task, which were modulated by learned material content and acquisition level before sleep (Maquet et al., 2000; Peigneux et al., 2003). While TMR is not often applied in REM, a number of studies have found significant effects (Hars et al., 1985; Guerrien et al., 1989; Smith and Weeden, 1990; Rihm and Rasch, 2015). Two studies of odour-based TMR have compared the impacts of TMR in REM and non-REM on a procedural memory task (Rasch et al., 2007; Laventure et al., 2016). While both of these found an impact of TMR in non-REM, neither showed an impact of REM TMR. Our findings are in keeping with these odour-based studies, since we found no behavioural benefits of REM. However, our own prior observation that the time spent in REM modulates neuroplasticity in the motor system as a result of TMR in SWS (Cousins et al., 2016) suggests that, even if it REM TMR does not directly benefit this task, REM does contribute to the consolidation of motor sequences.

Interestingly, our ERP results suggest that the brain can distinguish whether sounds are related to the left or right hand during REM. Indeed, previous research also suggests that discrimination of a stimulus’ significance and semantic content may persist during REM (Niiyama et al., 1994; Sallinen et al., 1996; Bastuji and García-Larrea, 1999; Takahara et al., 2006). However, these ERPs did not predict any form of consolidation which we were able to measure, and the fact that stimuli can be processed during REM does not mean that this processing will lead to behavioural impacts. It is of course possible that the type of plasticity mediated by REM simply does not impact upon the performance measures which we were able to collect. Thus, if we had collected fMRI data, we might have seen task-related plasticity in the REM TMR group.

### TMR preferentially benefits the non-dominant hand

Prior studies of how TMR impacts upon the serial reaction time task have typically used only the left (non-dominant) hand (Cousins et al., 2014, 2016; Schönauer et al., 2014). To determine whether TMR differentially impacts upon dominant and non-dominant hands, we modified the task such that an equal number of responses were required from each hand. Interestingly, we found that, while there is a trend towards a TMR effect in both hands, only responses in the non-dominant hand show a significant benefit.

Both raw RT and sequence-specific skill were worse in the left compared to the right hand. Our results thus largely fit with the idea that weaker memories particularly benefit from TMR (Drosopoulos et al., 2007; Cairney et al., 2016; Tambini et al., 2017; Schapiro et al., 2018). However, it may also be the case that some other aspect of the way processing in the dominant and non-dominant hand differs underpins the observed consolidation bias. Hand and finger movement representation in the primary motor cortex is significantly larger on the side contralateral to the dominant hand, which may be related to the greater motor skill often experienced in the preferred hand (Volkmann et al., 1998). It is possible that there is greater opportunity for gain in the non-dominant hand precisely because of its smaller motor skill repertoire. It is also possible that this result is related to the lateralised processing of hand movements rather than to a difference in handedness. The right motor cortex is primarily activated in response to contralateral (left) hand movements (Kim et al., 1993). In contrast, the left motor cortex is activated during both contralateral (right) and ipsilateral (left) hand movements, especially in right-handed participants. In other words, left-handed movements produce activity in both hemispheres (Gut et al., 2007), and it is possible that this more widespread activation could be at the basis of the left hand benefit which we find in our study. These ideas could be tested using a similar experiment with left handed participants. Such an experiment would help to clarify whether TMR is truly biased towards the non-dominant hand, or instead just towards the left hand.

Interestingly, the extent to which SWS TMR-elicited ERPs were greater in the experimental compared to the adaptation night was negatively associated with the extent of the LH cueing advantage the next day. This could be due to the phenomenon whereby more familiar sounds tend to elicit a weaker ERP (e.g. Niiyama et al., 1995; Cote and Campbell, 1999; Atienza et al., 2001). Thus, if the ERP in the experimental night is larger, it could indicate a lack of recognition, and a lack of association with the task and a correspondingly lower probability of eliciting reactivation. Alternatively, it is possible that this large early potential disrupts how much reactivation can take place, and if the brain is in a highly reactive state when TMR occurs then the manipulation is less effective. As a third possibility, the large ERP may indicate a weak sound-memory representation, and thus reflect a lot of effort (“I know this means something”) but resulting in little behavioural improvement.

## Conclusion

In the current study, we demonstrate that TMR in SWS, but not REM, leads to a consolidation benefit. Interestingly, our ERP results demonstrate that the brain does process TMR stimuli during REM, although such processing may occur in a manner that is not conducive to consolidation of our procedural task. We also show that the non-dominant hand benefits preferentially from TMR-cued memory consolidation. This may be because the non-dominant hand is weaker in the first place, allowing more space for improvement, or because this hand places more bilateral demands on the brain, and may thus draw on more of the neural circuity that benefits from TMR.

## Acknowledgments

This work was supported by the ERC grant SolutionSleep to PL and by a PhD studentship to ACMK from Cardiff University. ACMK, SB, and PL designed the study. ACMK, AM, MS, and TH collected the data and sleep scored the nights. ACMK, MEAA, and MR analysed the data. ACMK, MEAA, and PL wrote the manuscript. Our thanks go to James Cousins, Tia Tsimpanouli, Suliman Belal, and Monika Śledziowska for help with the task scripts. We also thank Miguel Navarrete and Matthias Treder for help with data analysis and advice on FieldTrip.

## Supplements

### Explicit Recall

For the explicit sequence knowledge scoring, individual items were only considered correct if they were in the correct position within the sequence, and if they were part of a segment that contains >2 correct items. This corresponded to the method used by Cousins et al. (2014).

To examine whether TMR leads to the overnight emergence of explicit knowledge, we conducted Wilcoxon signed-rank tests for each group separately. There was no difference between the cued and the uncued sequence in either the SWS or the REM group, see Table S1. In other words, TMR did not affect participants’ explicit knowledge of the sequences in this experiment.

**Table S1.** Explicit memory for each sequence, in number of correct items (± standard deviation).

|  | **REM sleep group**<br>*n* = 15 | **SWS group**<br>*n* = 16 |
| --- | --- | --- |
| **Cued sequence** | 10.1 ± 2.8 | 9.8 ± 2.9 |
| **Uncued sequence** | 10.7 ± 2.4 | 9.3 ± 3.5 |
| **Significance value** | *p* = 0.524 | *p* = 0.837 |

### Electrophysiology: Event-related potentials per night

Event-related potential (ERP) comparisons between the adaptation and experimental night were pre-processed and analysed in the same way as those concerning the left- and right-handed trials. In both the REM and the SWS group, ERPs to cues in the experimental night elicited a larger response than those in the adaptation night (see Figure S1a-b). However, this difference was not significant.

### Time-frequency analysis

Time-frequency analysis used a 5-cycle frequency-dependent Hanning taper to obtain spectral power from 4-30 Hz in frequency steps of 0.5 Hz and time steps of 5 ms. Averages across subject groups (REM or SWS) and cue groups (left-versus right-handed or adaptation versus experimental night) were calculated as power change relative to a baseline window of −1 second until cue onset. To capture both slow and fast changes in the time-frequency representations, we looked at the entire time interval from cue onset (time 0) until another tone was played approximately 1.5 seconds later. Statistical analyses of time-frequency comparisons between cues were performed as paired-samples t-tests and corrected for multiple comparisons with FieldTrip’s nonparametric cluster-based permutation method, using 1000 permutations. Results were considered significant at p < 0.05.

Time-frequency results comparing the nights in the REM group showed some slow activity approximately 250-500 ms after the cue (see Figure S1c-d). This activity seemed to occur in both nights, and a cluster permutation test confirmed that the difference between the nights was not significant. Results in the SWS group revealed early (around 500 ms after the cue) theta-band activity, and a weak fast spindle-band response to experimental cues around 1.2 seconds after cue onset (see Figure S1e-f). Again, these responses were similar between the nights, and statistical analyses showed that the difference was not significant. When looking at the left- and right-handed cues during the experimental night, we found that the REM group again showed early theta or even delta activity at around 250-500 ms after the cue (see Figure S2a-b). In response to the left-handed cues there also seemed to be some fast spindle-band activity starting around 500 ms after the cue, while this was not the case when looking at the right-handed cues. However, statistical analyses did not show a significant difference between the EEG responses to these different cues. This remained the case when the left and right hemisphere were analysed separately.

In the SWS group, both time-frequency results showed theta-band activity around 500 ms after the cue (see Figure S2c-d). This activity even reached the alpha band in response to right-handed cues. In contrast, there seemed to be a slightly more pronounced spindle-band response around 1.1-1.2 seconds after left-handed cues. Statistical analyses comparing these responses did not reveal any significant differences. The two hemispheres showed remarkably similar results, and differences in the time-frequency responses to left- and right-handed cues thus remained non-significant when we analysed the hemispheres separately.

**Figure S1.**
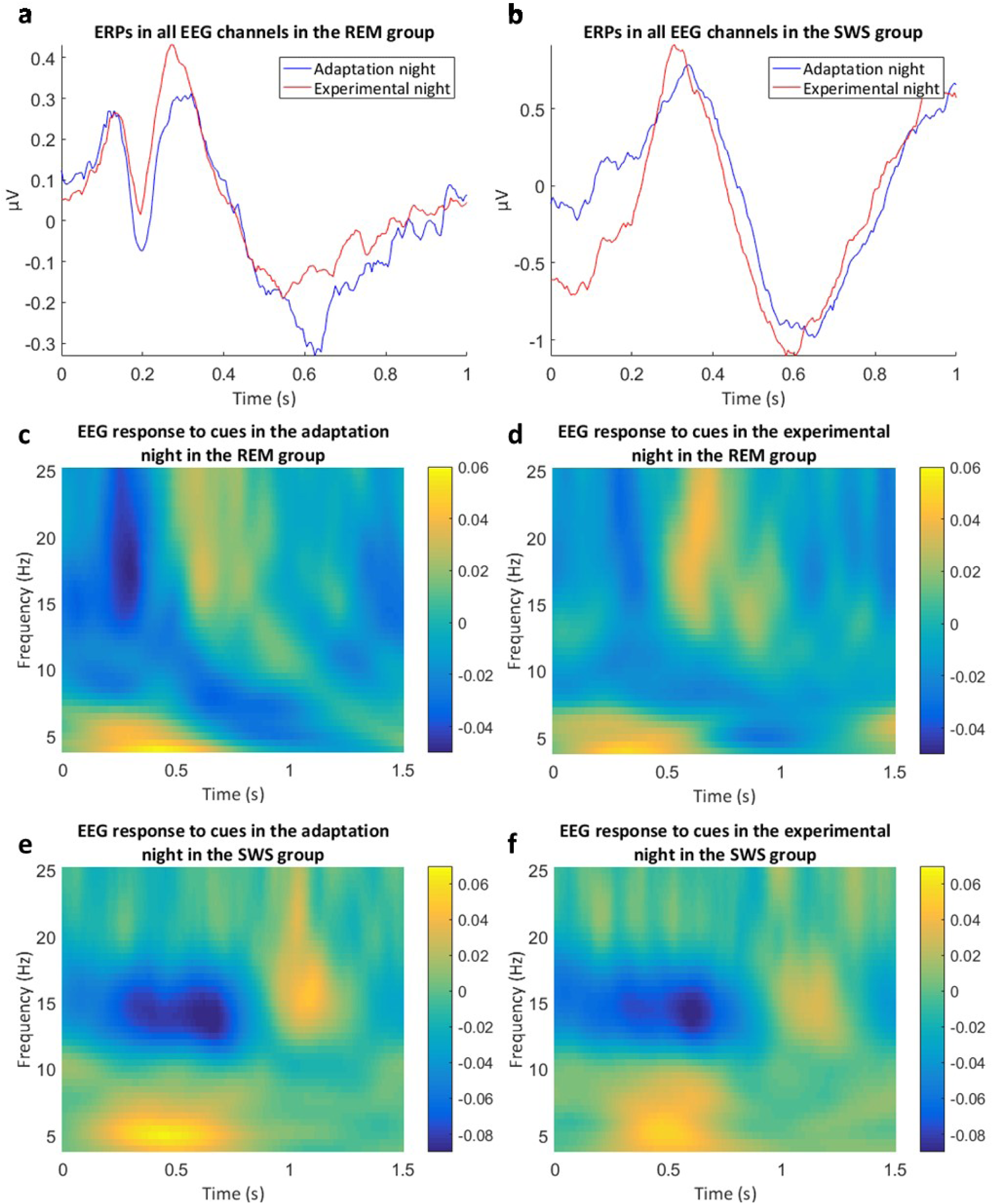
Electrophysiological results comparing the adaptation and experimental night in all EEG channels. **a)** ERPs in all EEG channels in the REM group. **b)** ERPs in all EEG channels in the SWS group. **c)** Time-frequency representation of the EEG response to cues in the adaptation night in the REM group. **d)** Time-frequency representation of the EEG response to cues in the experimental night in the REM group. **e)** Time-frequency representation of the EEG response to cues in the adaptation night in the SWS group. **f)** Time-frequency representation of the EEG response to cues in the experimental night in the SWS group. Legends in the time-frequency plots represent relative change from baseline.

**Figure S2.**
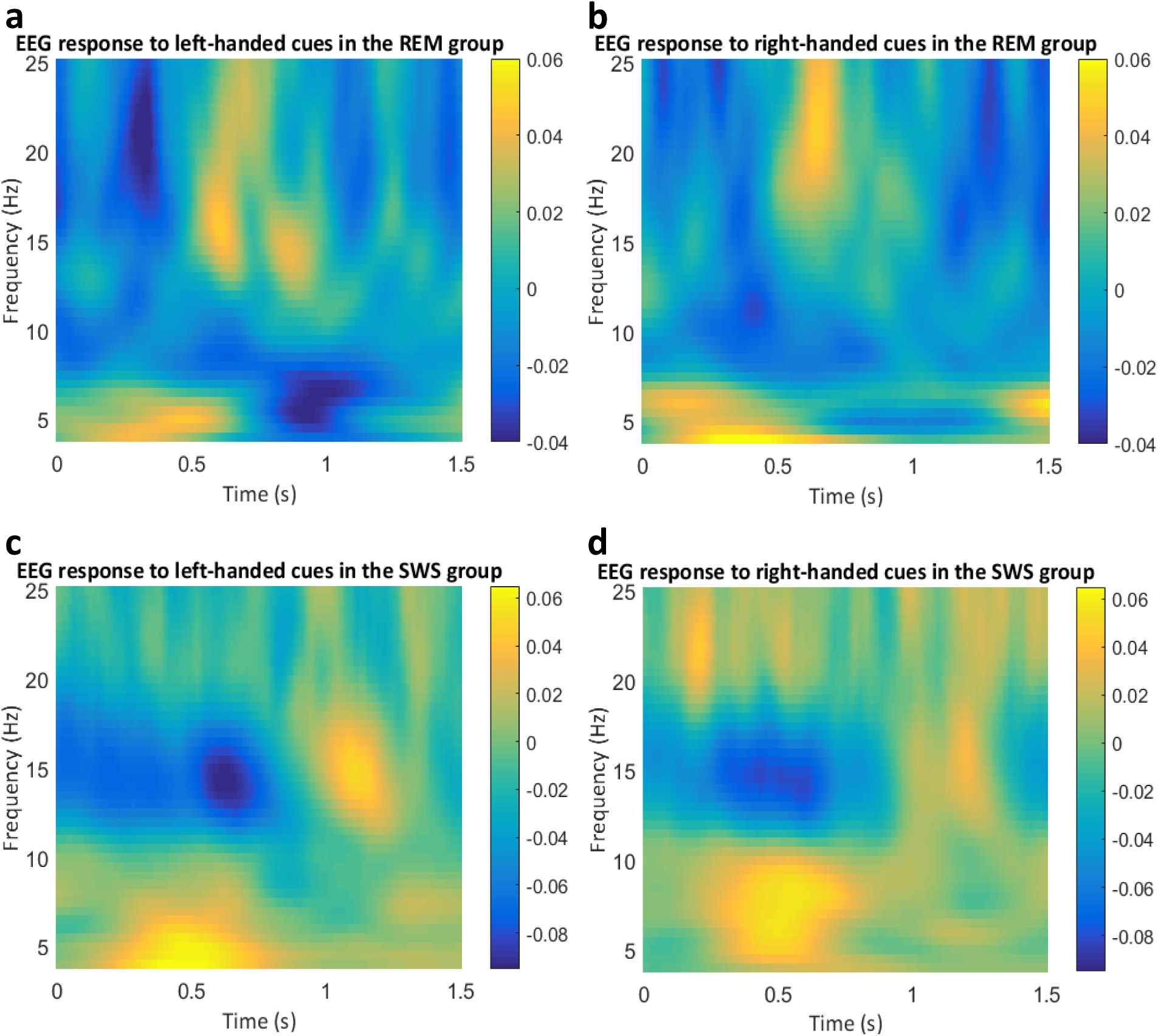
Time-frequency results comparing cues related to the left and right hand, in all EEG channels. Legends represent relative change from baseline. **a)** Time-frequency representation of the EEG response to cues associated with the left hand in the REM group. b) Time-frequency representation of the EEG response to cues associated with the right hand in the REM group. **c)** Time-frequency representation of the EEG response to cues associated with the left hand in the SWS group. d) Time-frequency representation of the EEG response to cues associated with the right hand in the SWS group.

## References

Åkerstedt T, Gillberg M (1990) Subjective and Objective Sleepiness in the Active Individual. Int J Neurosci 52:29–37.

Antony JW, Gobel EW, O’Hare JK, Reber PJ, Paller KA (2012) Cued memory reactivation during sleep influences skill learning. Nat Neurosci 15:1114–1116.

Antony JW, Piloto L, Wang M, Pacheco P, Norman KA, Paller KA (2018) Sleep Spindle Refractoriness Segregates Periods of Memory Reactivation. Curr Biol 28:1736-1743.e4.

Atienza M, Cantero JL, Escera C (2001) Auditory information processing during human sleep as revealed by event-related brain potentials. Clin Neurophysiol 112:2031–2045.

Bakeman R (2005) Recommended effect size statistics for repeated measures designs. Behav Res Methods 37:379–384.

Bastuji H, García-Larrea L (1999) Evoked potentials as a tool for the investigation of human sleep. Sleep Med Rev 3:23–45.

Belal S, Cousins JN, El-Deredy W, Parkes L, Schneider J, Tsujimura H, Zoumpoulaki A, Perapoch M, Santamaria L, Lewis P (2018) Identification of memory reactivation during sleep by EEG classification. Neuroimage 176:203–214.

Berry RB, Brooks R, Gamaldo CE, Harding SM, Lloyd RM, Marcus CL, Vaughn B V (2015) The AASM Manual for the Scoring of Sleep and Associated Events. Rules, terminology and technical specifications. Darien, IL: American Academy of Sleep Medicine.

Booth V, Poe GR (2006) Input source and strength influences overall firing phase of model hippocampal CA1 pyramidal cells during theta: Relevance to REM sleep reactivation and memory consolidation. Hippocampus 16:161–173.

Buysse DJ, Reynolds CF, Monk TH, Berman SR, Kupfer DJ (1989) The Pittsburgh Sleep Quality Index (PSQI): A new instrument for psychiatric research and practice. Psychiatry Res 28:193–213.

Cairney SA, Durrant SJ, Hulleman J, Lewis PA (2014) Targeted Memory Reactivation During Slow Wave Sleep Facilitates Emotional Memory Consolidation. Sleep 37:701–707.

Cairney SA, Lindsay S, Sobczak JM, Paller KA, Gaskell MG (2016) The Benefits of Targeted Memory Reactivation for Consolidation in Sleep are Contingent on Memory Accuracy and Direct Cue-Memory Associations. Sleep 39:1139– 1150.

Cellini N, Cappuzo A (2018) Shaping memory consolidation via targeted memory. Ann N Y Acad Sci 1426:52–71.

Cote KA, Campbell KB (1999) P300 to high intensity stimuli during REM sleep. Clin Neurophysiol 110:1345–1350.

Cousins JN, El-Deredy W, Parkes LM, Hennies N, Lewis PA (2014) Cued Memory Reactivation during Slow-Wave Sleep Promotes Explicit Knowledge of a Motor Sequence. J Neurosci 34:15870–15876.

Cousins JN, El-Deredy W, Parkes LM, Hennies N, Lewis PA (2016) Cued Reactivation of Motor Learning during Sleep Leads to Overnight Changes in Functional Brain Activity and Connectivity. PLOS Biol 14:e1002451.

Diekelmann S, Born J (2010) The memory function of sleep. Nat Rev Neurosci 11:114–126.

Drosopoulos S, Schulze C, Fischer S, Born J (2007) Sleep’s function in the spontaneous recovery and consolidation of memories. J Exp Psychol Gen 136:169–183.

Fritz CO, Morris PE, Richler JJ (2012) Effect size estimates: Current use, calculations, and interpretation. J Exp Psychol Gen 141:2–18.

Fuentemilla L, Miró J, Ripollés P, Vilà-Balló A, Juncadella M, Castañer S, Salord N, Monasterio C, Falip M, Rodríguez-Fornells A (2013) Hippocampus-Dependent Strengthening of Targeted Memories via Reactivation during Sleep in Humans. Curr Biol 23:1769–1775.

Görtelmeyer R (1985) On the development of a standardized sleep inventory for the assessment of sleep. In: Methods of Sleep Research (Kubicki S, Herrmann WM, eds), pp 93–98. Stuttgart: Gustav Fischer.

Guerrien A, Dujardin K, Mandai O, Sockeel P, Leconte P (1989) Enhancement of memory by auditory stimulation during postlearning REM sleep in humans. Physiol Behav 45:947–950.

Gut M, Urbanik A, Forsberg L, Binder M, Rymarczyk K, Sobiecka B, Kozub J, Grabowska A (2007) Brain correlates of right-handedness. Acta Neurobiol Exp (Wars) 67:43–51.

Hars B, Hennevin E, Pasques P (1985) Improvement of learning by cueing during postlearning paradoxical sleep. Behav Brain Res 18:241–250.

Hauner KK, Howard JD, Zelano C, Gottfried JA (2013) Stimulus-specific enhancement of fear extinction during slow-wave sleep. Nat Neurosci 16:1553–1555.

Hoddes E, Zarcone V, Smythe H, Phillips R, Dement WC (1973) Quantification of sleepiness: a new approach. Psychophysiology 10:431–436.

Howe T, Wilson MA, Ji D, Jones MW (2019) Extending evidence for REM-associated replay in hippocampal CA1 place cells. In: Program No. 333.10.2019 Neuroscience Meeting Planner, pp Online. Chicago, IL: Society for Neuroscience. Available at: https://www.abstractsonline.com/pp8/#!/7883/presentation/67466.

Karni A, Tanne D, Rubenstein BS, Askenasy JJM, Sagi D (1994) Dependence on REM sleep of overnight improvement of a perceptual skill. Science 265:679–682.

Kim SG, Ashe J, Hendrich K, Ellermann JM, Merkle H, Uǧurbil K, Georgopoulos AP (1993) Functional magnetic resonance imaging of motor cortex: Hemispheric asymmetry and handedness. Science 261:615–617.

Korman M, Doyon J, Doljansky J, Carrier J, Dagan Y, Karni A (2007) Daytime sleep condenses the time course of motor memory consolidation. Nat Neurosci 10:1206–1213.

Korman M, Raz N, Flash T, Karni A (2003) Multiple shifts in the representation of a motor sequence during the acquisition of skilled performance. Proc Natl Acad Sci U S A 100:12492–12497.

Lakens D (2013) Calculating and reporting effect sizes to facilitate cumulative science: a practical primer for t-tests and ANOVAs. Front Psychol 4:863.

Laventure S, Fogel S, Lungu O, Albouy G, Sévigny-Dupont P, Vien C, Sayour C, Carrier J, Benali H, Doyon J (2016) NREM2 and Sleep Spindles Are Instrumental to the Consolidation of Motor Sequence Memories. PLoS Biol 14:1–27.

Louie K, Wilson MA (2001) Temporally structured replay of awake hippocampal ensemble activity during rapid eye movement sleep. Neuron 29:145–156.

Mangiafico S (2020) rcompanion: Functions to Support Extension Education Program Evaluation.: R package version 2.3.25 Available at: https://cran.r-project.org/package=rcompanion.

Maquet P, Laureys S, Peigneux P, Fuchs S, Petiau C, Phillips C, Aerts J, Del Fiore G, Degueldre C, Meulemans T, Luxen A, Franck G, Van Der Linden M, Smith C, Cleeremans A (2000) Experience-dependent changes in changes in cerebral activation during human REM sleep. Nat Neurosci 3:831–836.

Maquet P, Schwartz S, Passingham R, Frith C (2003) Sleep-related consolidation of a visuomotor skill: Brain mechanisms as assessed by functional magnetic resonance imaging. J Neurosci 23:1432–1440.

Navarrete M, Schneider J, Ngo H-V V, Valderrama M, Casson AJ, Lewis PA (2020) Examining the optimal timing for closed-loop auditory stimulation of slow-wave sleep in young and older adults. Sleep 43:zsz315.

Niiyama Y, Fujiwara R, Satoh N, Hishikawa Y (1994) Endogenous components of event-related potential appearing during NREM stage 1 and REM sleep in man. Int J Psychophysiol 17:165–174.

Niiyama Y, Fushimi M, Sekine A, Hishikawa Y (1995) K-complex evoked in NREM sleep is accompanied by a slow negative potential related to cognitive process. Electroencephalogr Clin Neurophysiol 95:27– 33.

Nishida M, Walker MP (2007) Daytime Naps, Motor Memory Consolidation and Regionally Specific Sleep Spindles. PLoS One 2:e341.

Olejnik S, Algina J (2003) Generalized Eta and Omega Squared Statistics: Measures of Effect Size for Some Common Research Designs. Psychol Methods 8:434–447.

Oostenveld R, Fries P, Maris E, Schoffelen J-M (2011) FieldTrip: Open source software for advanced analysis of MEG, EEG, and invasive electrophysiological data. Comput Intell Neurosci 2011:Article ID: 156869.

Peigneux P, Laureys S, Fuchs S, Destrebecqz A, Collette F, Delbeuck X, Phillips C, Aerts J, Del Fiore G, Degueldre C, Luxen A, Cleeremans A, Maquet P (2003) Learned material content and acquisition level modulate cerebral reactivation during posttraining rapid-eye-movements sleep. Neuroimage 20:125–134.

Poe GR, Nitz DA, McNaughton BL, Barnes CA (2000) Experience-dependent phase-reversal of hippocampal neuron firing during REM sleep. Brain Res 855:176–180.

Purcell SM, Manoach DS, Demanuele C, Cade BE, Mariani S, Cox R, Panagiotaropoulou G, Saxena R, Pan JQ, Smoller JW, Redline S, Stickgold R (2017) Characterizing sleep spindles in 11,630 individuals from the National Sleep Research Resource. Nat Commun 8:15930.

R Core Team (2020) R: A language and environment for statistical computing. Available at: https://www.r-project.org/.

Rasch B, Born J (2013) About sleep’s role in memory. Physiol Rev 93:681–766.

Rasch B, Büchel C, Gais S, Born J (2007) Odor cues during Slow-Wave Sleep prompt declarative memory consolidation. Science 315:1426– 1429.

Ridding MC, Flavel SC (2006) Induction of plasticity in the dominant and non-dominant motor cortices of humans. Exp Brain Res 171:551– 557.

Rihm JS, Rasch B (2015) Replay of conditioned stimuli during late REM and stage N2 sleep influences affective tone rather than emotional memory strength. Neurobiol Learn Mem 122:142–151.

Rudoy JD, Voss JL, Westerberg CE, Paller KA (2009) Strengthening Individual Memories by Reactivating Them During Sleep. Science 326:1079.

Sallinen M, Kaartinen J, Lyytinen H (1996) Processing of auditory stimuli during tonic and phasic periods of REM sleep as revealed by event-related brain potentials. J Sleep Res 5:220–228.

Schapiro AC, McDevitt EA, Rogers TT, Mednick SC, Norman KA (2018) Human hippocampal replay during rest prioritizes weakly learned information and predicts memory performance. Nat Commun 9:3920.

Schönauer M, Alizadeh S, Jamalabadi H, Abraham A, Pawlizki A, Gais S (2017) Decoding material-specific memory reprocessing during sleep in humans. Nat Commun 8:15404.

Schönauer M, Geisler T, Gais S (2014) Strengthening Procedural Memories by Reactivation in Sleep. J Cogn Neurosci 26:143–153.

Schreiner T, Doeller CF, Jensen O, Rasch B, Staudigl T (2018) Theta Phase-Coordinated Memory Reactivation Reoccurs in a Slow-Oscillatory Rhythm during NREM Sleep. Cell Rep 25:296– 301.

Shanahan LK, Gjorgieva E, Paller KA, Kahnt T, Gottfried JA (2018) Odor-evoked category reactivation in human ventromedial prefrontal cortex during sleep promotes memory consolidation. Elife 7:e39681.

Singmann H, Bolker B, Westfall J, Aust F, Ben-Shachar MS (2020) afex: Analysis of Factorial Experiments.: R package version 0.26–0 Available at: https://cran.r-project.org/package=afex.

Skaggs WE, McNaughton BL (1996) Replay of Neuronal Firing Sequences in Rat Hippocampus During Sleep Following Spatial Experience. Science 271:1870–1873.

Smith C (1993) REM Sleep and Learning: Some Recent Findings. In: The Functions of Dreaming (Moffitt A, Kramer M, Hoffmann R, eds), pp 341–361. Albany, NY: State University of New York Press.

Smith C (1995) Sleep states and memory processes. Behav Brain Res 69:137–145.

Smith C (2001) Sleep states and memory processes in humans: Procedural versus declarative memory systems. Sleep Med Rev 5:491–506.

Smith C, Smith D (2003) Ingestion of Ethanol Just Prior to Sleep Onset Impairs Memory for Procedural but not Declarative Tasks. Sleep 26:185–191.

Smith C, Weeden K (1990) Post training REMs coincident auditory stimulation enhances memory in humans. Psychiatr J Univ Ottawa 15:85–90.

Spencer RMC, Sunm M, Ivry RB (2006) Sleep-Dependent Consolidation of Contextual Learning. Curr Biol 16:1001–1005.

Takahara M, Nittono H, Hori T (2006) Effect of Voluntary Attention on Auditory Processing During REM Sleep. Sleep 29:975–982.

Tambini A, Berners-Lee A, Davachi L (2017) Brief targeted memory reactivation during the awake state enhances memory stability and benefits the weakest memories. Sci Rep 7:15325.

Veale JF (2014) Edinburgh Handedness Inventory - Short Form: A revised version based on confirmatory factor analysis. Laterality Asymmetries Body, Brain Cogn 19:164–177.

Volkmann J, Schnitzler A, Witte OW, Freund H-J (1998) Handedness and Asymmetry of Hand Representation in Human Motor Cortex. J Neurophysiol 79:2149–2154.

Walker MP, Brakefield T, Hobson JA, Stickgold R (2003a) Dissociable stages of human memory consolidation and reconsolidation. Nature 425:616–620.

Walker MP, Brakefield T, Morgan A, Hobson JA, Stickgold R (2002) Practice with sleep makes perfect: Sleep-dependent motor skill learning. Neuron 35:205–211.

Walker MP, Brakefield T, Seidman J, Morgan A, Hobson JA, Stickgold R (2003b) Sleep and the time course of motor skill learning. Learn Mem 10:275–284.

Walker MP, Stickgold R, Alsop D, Gaab N, Schlaug G (2005) Sleep-dependent motor memory plasticity in the human brain. Neuroscience 133:911–917.

Williams SE, Cumming J, Ntoumanis N, Nordin-Bates SM, Ramsey R, Hall C (2012) Further validation and development of the Movement Imagery Questionnaire. J Sport Exerc Psychol 34:621–646.

Wilson MA, McNaughton BL (1994) Reactivation of Hippocampal Ensemble Memories During Sleep. Science 265:676–679.

